# TGF-β1 enhances mouse mast cell release of IL-6 and IL-13

**DOI:** 10.1101/220665

**Authors:** David O. Lyons, Michele R. Plewes, Nicholas A. Pullen

## Abstract

For immune cells, TGF-β1 can enhance or repress effector functions. Here, we characterize the effects of TGF-β1 on IgE-mediated activation of primary murine mast cells derived from hematopoietic stem cells (BMMC). We also investigated potential interaction between TGF-β1 and stem-cell factor (SCF). Resting IL-6 production was increased with TGF-β1 but significance was lost following BMMC activation via IgE receptor (FcεRI) crosslinking. SCF also enhanced resting levels of IL-6, but there was no difference from control once FcεRI was engaged. SCF had no effect on IL-13 production; however, TGF-β1 treatment enhanced release of IL-13 upon FcεRI activation. Lastly, percent colocalization of SCF receptor (CD117) and FcεRI were unaffected by TGF-β1 treatment. These data reveal a novel positive effect of soluble TGF-β1 on mast cell activation.

## Introduction

Transforming Growth Factor Beta (TGF-β1) is a widely expressed cytokine. The TGF-ß1 signaling pathway evolved approximately one billion years ago as an immune regulatory mechanism among vertebrates. TGF-ß1 modulates cellular responses starting with binding to TGF-β receptor II (TGF-ßRII). TGF-ßRII then aggregates with TGF-βRI at the cell surface inducing the phosphorylation of Smad proteins intracellularly. Ultimately, this cascade reaches the nucleus for transcriptional regulation. TGF-ß1 is documented as generally inhibitory starting in the 1990s: as an anti-inflammatory, anti-autoimmune cytokine [1].

Mast cells are myeloid lineage cells of hematopoietic origin. They are present in the skin and along mucosal membranes, especially the gut where they combat helminth parasites. Well known for their roles in allergic pathologies, mast cells also are key to physiology at mucosal barriers. Mast cells are capable of collecting and presenting antigen to other cells [2]. They are also central in driving a T_H_2 response inducing B cells to class switch to IgE via IL-4 and IL-13 [3]. Canonical activation of mast cells starts with the priming of their high affinity IgE receptor, FcϵRI. In the body, mast cells are stably coated with IgE bound to FcϵRI. Upon multivalent antigen binding of these IgE-FcϵRI complexes, the receptors cluster – crosslink, and internalize, triggering signaling cascades resulting in degranulation of the cell and activation of transcription factors, such as STAT5, to upregulate cytokine production [3]. Hours to days later these cytokines (*e.g.*, IL-6, IL-13) are secreted [2]. They also exhibit an alternative pathway to activation through IL-33 and its receptor (ST2), independent of FcϵRI [4].

Recent evidence suggests that the interaction between mast cells and typical immunosuppressive cytokines varies from other immune cells. A typical inhibitory cytokine, IL-10, was shown to behave as an immunostimulant when given to mast cells and in the development of mucosal food allergies [5]. At the post-transcriptional level, IL-10 was found to regulate microRNAs, which enhanced skin mast cell secretion of IL-6 and IL-13 [6]. These interactions necessitate closer examination of the behavior of mast cells regarding common stimulatory and inhibitory molecules. Herein, we describe effects of TGF-ß1 on IgE-mediated mast cell activation by observing the production of two cytokines commonly associated with this process: IL-6 (an inflammatory T_H_-1/T_H_-17 cytokine) and IL-13 (a chemotactic cytokine that drives a T_H_-2 response). We demonstrate that soluble TGF-ß1 amplifies mast cell release of these cytokines. This effect opposes SCF exposure, depending on the cytokine; and it is unlikely due to receptor interference at the membrane.

## Methods

### Mice

C57/BL6 mice were housed and humanely euthanized according to University of Northern Colorado IACUC protocol #1702C-NP-M-20. When possible, tissue was obtained from control mice scheduled for other experiments.

### BMMC Differentiation

BMMC were cultured in RPMI 1640 (Thermo Fisher) supplemented with: 10% FBS (VWR), 1% pen/strep, 2mM L-Glutamine, 1mM Sodium Pyruvate, 10mM HEPES (Thermo Fisher), 30ng/mL IL-3 (Peprotech), and 0.05mM 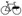-mercaptoethanol (Bio-Rad). Bone marrow was flushed from mouse femurs and tibias, then cleared of erythrocytes with ACK lysis buffer (Quality Biological). Cultures were initiated at 500,000 cells/mL in T75 flasks (VWR); individual mouse cultures were kept separate. Cultures were observed daily, if adherent cells were present, desired suspension cells were transferred to a new flask. Cultures were maintained between 250,000 and 1,000,000 cells/mL for 4-6 weeks to achieve pure BMMCs. All cultures were randomly divided into SCF-treated and untreated groups during differentiation. SCF-treated cell populations were given 10ng/mL SCF (Peprotech).

### TGF-ß1 Treatment

For TGF-ß1 treated groups, SCF-treated and untreated cells were distributed in 6-well plates at a density between 250,000 and 1,000,000 cells/mL. Cells were treated with 2ng/mL murine TGF-ß1 (Cell Signaling) and maintained for two days before IgE priming.

### IgE-mediated activation and quantification of BMMC cytokines

Cells were incubated overnight with 500ng/mL of IgE (BD Biosciences; 557079) specific to trinitrophenyl keyhole limpet hemocyanin (TNP-KLH; Santa Cruz Biotechnologies). Treatments were rinsed with RPMI and re-suspended in complete medium. The IgE-FcϵRI complexes were crosslinked by adding 300ng/mL TNP-KLH and incubating overnight (IgE-exposed, no TNP-KLH controls for each group included). ELISAs were performed on conditioned media according to the manufacturer’s protocols for IL-6 (PeproTech; 900-T50) and IL-13 (PeproTech; 900-K207).

### Microscopy

Cells were treated (TFG-β or IgE-XL) as stated above. Following treatment, cells were rinsed with 1× PBS, fixed (4% PFA), and blocked (5% normal goat serum).

Primary antibodies (FITC-CD117 (mouse-mAB; 1:50; Biolegend #105805), Alexa647 FcϵR1 (mouse-mAB; 1:50; Biolegend #134309), and TGFβRII (rabbit-mAB; 1:50; ABClonal #A11765)) were incubated at 4°C for 24 h. Following incubation, cells were rinsed and anti-rabbit Alexa568 (1:500; Life Technologies #A11011) was incubated for 1h at room temperature. Cells were mounted onto microscope slide using 10µL SlowFade-Gold DAPI mounting medium (Life Technologies #S36938) and No. 1 cover slip. Images were collected using a Zeiss confocal microscope equipped with a 100× oil objective (N.A. = 1.4), acquisition image size 512×512 pixels (33.3µm×33.3µm), light collection 450nm-1000nm. Thirty cells were randomly selected from each slide for analysis. The JACoP plug-in for ImageJ was used to determine the Manders’ coefficient for each image. Coefficients ranged from 0 to 1, which were then transformed into percent colocalization by multiplying by 100.

### Statistics

Cytokine secretion was measured in pg/mL per 10^6^ cells. Cytokine levels were also quantified based on the fold change compared to the untreated controls. Data was tested for normality using the Shapiro-Wilks test and transformed accordingly.

Cytokine release data were analyzed via Welch’s t-test. Significance was noted at p≤0.05. Fluoresence microscopy data are reported as least square means±standard error of the mean with significance at p<0.05. The effects of TGF-ß1 and FcϵRI crosslinking on colocalization of BMMC surface receptors were analyzed using one-way analysis of variance.

## Results

### TGF-ß1 enhances IL-6 secretion by BMMCs

BMMCs were treated with TGF-ß1, SCF, or both, and then activated by crosslinking FcϵRI. The following IL-6 secretion data were collected from at least 4 separate BMMC populations (*i.e.*, bone marrow differentiated from different mice). IL-6 was increased following IgE-mediated activation (Figure 1 a), but IL-6 was also increased in cells treated with either SCF or TGF-ß1 alone independent of IgE-mediated activation; cells treated with both ligands exhibited higher levels compared to untreated controls (Figure 1 c). When cross-linked, only TGF-ß1 appeared to enhance IL-6 secretion but the difference was not significant (Data not shown). These data show that soluble TGF-ß1 directly enhances mast cell IL-6 production independent of IgE-mediated activation.

**Figure 1:**
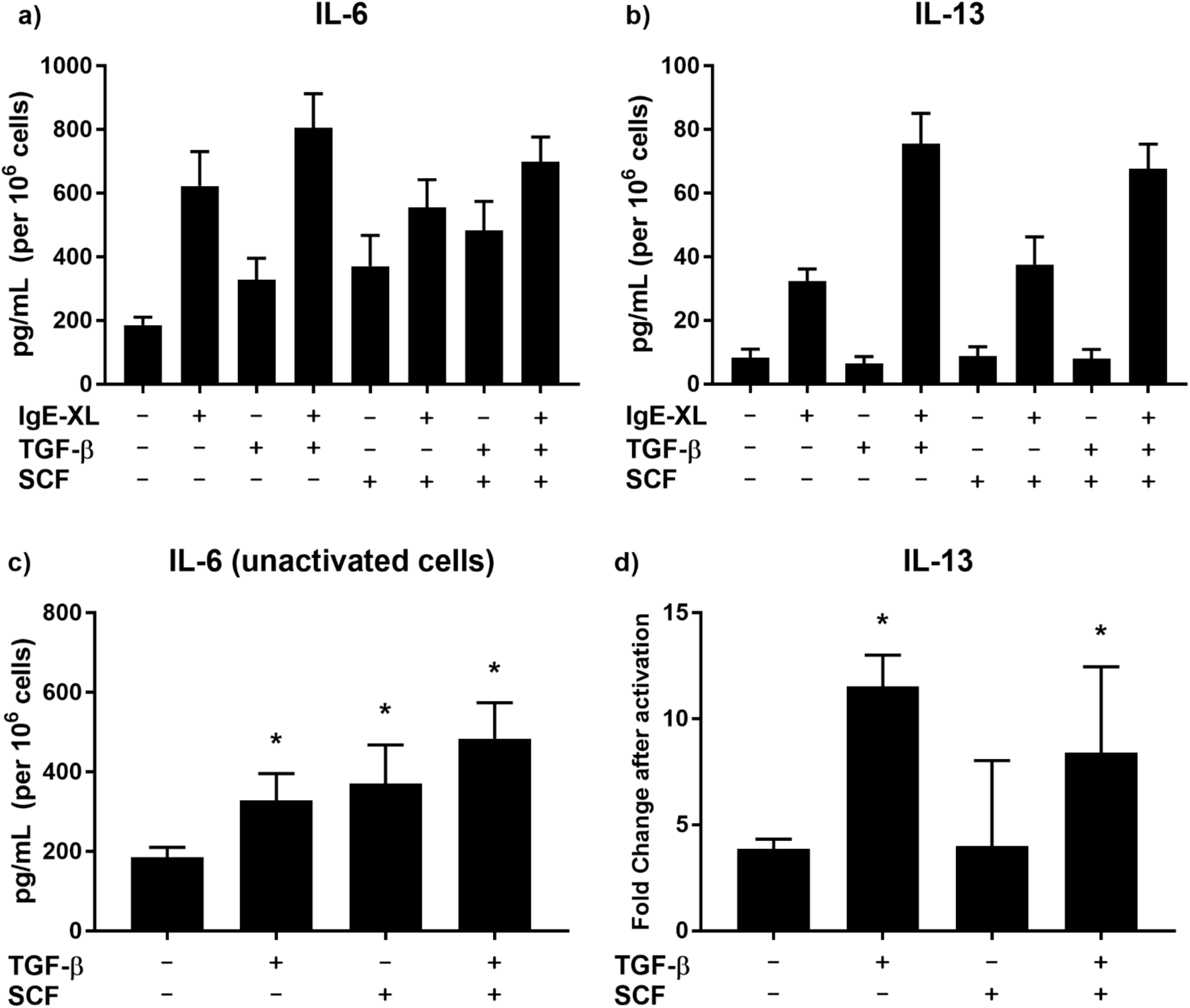
IL-6 and IL-13 secretion by mast cells based on treatment. **(a and b)** Raw cytokine production for IL-6 **(a)** and IL-13 **(b)** corrected for the number of cells assayed. **(c)** IL-6 production in resting (un-crosslinked) BMMCs. **(d)** Fold change in IL-13 production following mast cell activation.

### TGF-ß1, but not SCF, enhances IL-13 secretion from BMMCs

Concentrations of IL-13 were also measured as the fold change when crosslinked. Un-crosslinked populations secreted little to no detectable IL-13. Figure 1 b shows the raw IL-13 production based on treatment. Unlike IL-6, IL-13 production prior to IgE activation was not affected based on cytokine treatment. However, treatment with TGF-ß1 resulted in higher levels of IL-13 upon IgE-mediated activation – this effect persisted when co-treated with SCF, but not with SCF alone (Figure 1 d). These data show that soluble TGF-ß1 enhances IL-13 production upon FcϵRI engagement.

### Crosslinking decreases FcϵRI‐ CD117 (c-kit) percent colocalization, independent of TGF-ß1 treatment

Percent colocalization of FcϵRI-CD117, TGF-βRII-CD117, and TGF-βRII-FcϵRI were dertermined using immunofluorescence microscopy (Figure 2). Crosslinking (+XL) BMMCs decreased percent colocalization of FcϵRI and CD117 in the presence and absence of TFG-ß (Figure 2 b). Additionally, the decreased percent colocalization was proportional between treatment groups, indicating TGF-β1 does not directly affect surface receptor function on mast cells (Figure 2 c).

**Figure 2:**
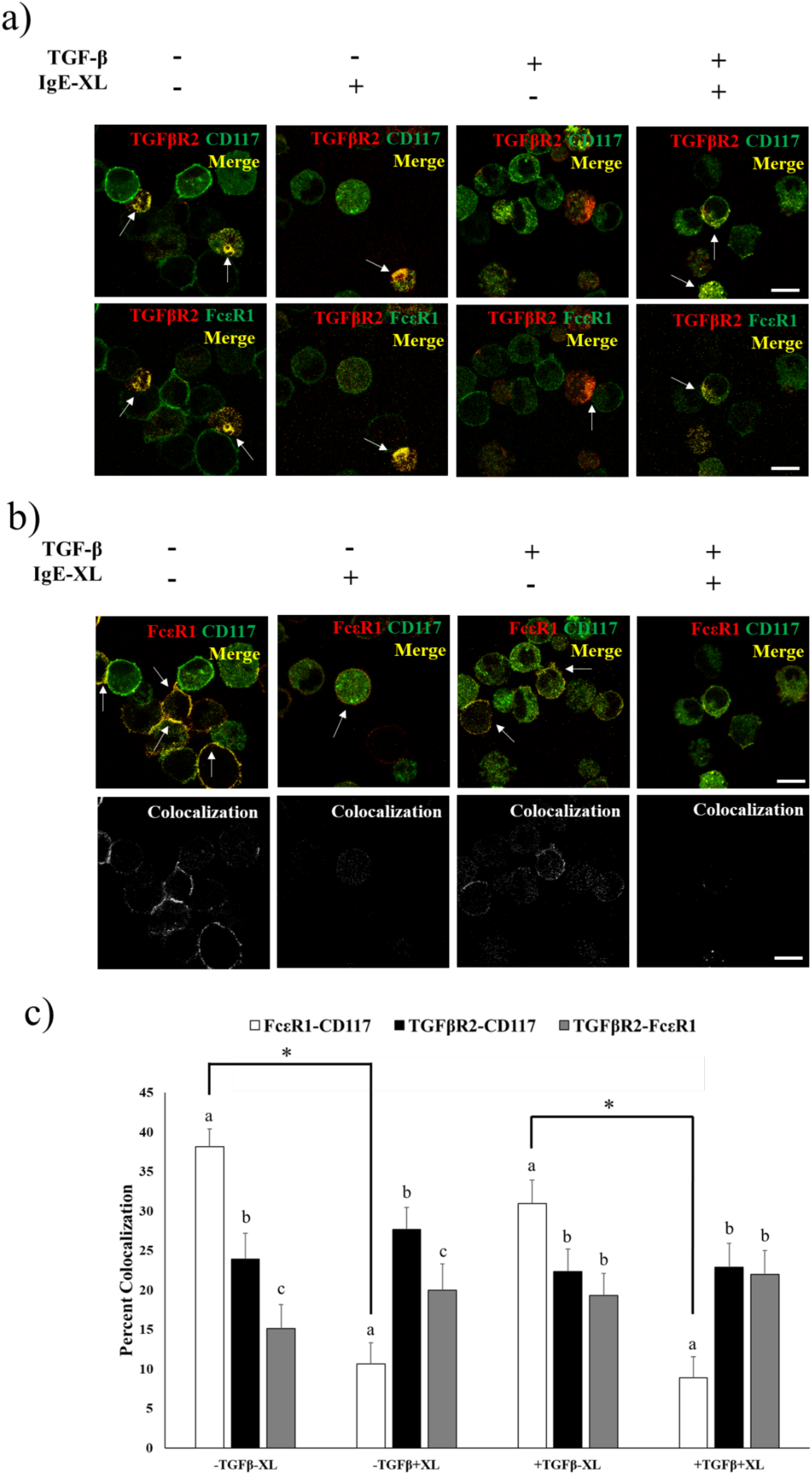
Colocalization of TGF-βRII, FcϵRI and CD117. **(a)** Representative micrographs of TFG-ßRII with FcϵRI or CD117. **(b)** Representative micrographs of FcϵRI and CD117; corresponding colocalization shown below. **(c)** Quantitative analysis of colocalization within treatment. Arrow represent regions of colocalization. Micron bar represents 10µm. Means with different letters^abc^ differ significantly within treatment (P < 0.05).

## Discussion

In our experiments soluble TGF-ß1 stimulates IL-6 secretion independent of IgE-mediated activation. TGF-ß1 has been shown to promote mast cell IL-6 production in the context of lung inflammation; this promotes neutrophil apoptosis and clearance [7]. However, this mechanism involved T_reg_ cell surface sequestered TGF-ß1 [8]. Our findings suggest that TGF-ß1 plays a directly stimulatory role on IL-6 production, potentially independent of immunosuppressive T_reg_, which could elicit acute phase inflammatory responses. Characterizing the mechanism(s) involved and carefully testing the direct effect(s) of soluble TGF-ß1 on other myeloid cells is warranted. Alternatively, myeloid-derived suppressor cells (MDSC) utilize IL-6 to support tumor progression [9]. MDSCs also enhance IL-6 and IL-13 secretion by activated mast cells [10]. The careful study of interactions between TGF-ß1, IL-6, T_reg_, and mast cells will be insightful in chronic inflammatory settings, such as high-grade solid cancers. Here, TGF-ß1 also enhances production of IL-13 following mast cell activation via FcϵRI. IL-13 is a major cytokine in T_H_2-related immune responses, likey responsible for the clearing of large extracellular insults, such as gut parasites [11]. However, unmitigated IL-13 release will drive B-lymphocyte class switching to IgE, which in turn coats naïve mast cells via FcϵRI, thus prompting a vicious T_H_2 cycle [12]. Our experiments show that TGF-ß1 directly enhances IL-13 production, which propagates such pathologies.

Through fluorescence microscopy we observed any effects TGF-ß1, and its receptor TGFBRII, might have at the signaling apex of the definitive mast cell surface receptors FcϵRI and CD117 (SCF receptor). FcϵRI and CD117 colocalization is lost upon crosslinking of FcϵRI with IgE-antigen complexes. This is not surprising as FcϵRI internalization after cross-linking is known [13]. However, this decline in colocalization is unchanged with TGF-ß1 treatment, suggesting there is no apparent surface cross-talk occurring among these receptors. It has been shown that TGF-ß1 transcriptionally represses FcϵRI and CD117 through regulation of Etf homologous factor (Ehf) [14]. This inhibition was not observed in our studies, suggesting reduced receptor expression of CD117 through flow cytometric analysis does not directly correspond to reduced receptor expression.

We demonstrate that TGF-ß1 plays a direct stimulatory role on primary mast cells *in vitro*. The specific molecular mechanism underlying this effect is still unknown. TGF-ß1 might modulate targets outside of the canonical TGF-ß1 signaling cascade (Smads) as preliminary experiments using a TGF-ßR1 inhibitor resulted in no significant differences in cytokine expression (data not shown). IL-6 secretion was increased independent of IgE-mediated activation, suggesting that TGF-ß1 non-canonically targets the MAP kinase or Akt pathways to enhance IL-6 production [15]. Future research into the phosphorylation patterns of this pathway will reveal the specific effects of TGF-ß1. Additionally, IL-13 production was altered only after IgE activation, suggesting that TGF-ß1 modulates the FcϵRI signaling pathway *(e.g*., STAT5). Altogether these data necessitate careful examination of soluble TGF-ß1 with respect to mast cell effector functions.

## Authors’ Contributions

DL undertook experiments, design and data analysis, and drafted the manuscript; MP performed microscopic analyses; NP conceived, designed, and coordinated the study, and assisted manuscript drafting. All authors gave final approval for publication.

## Competing Interests

The authors declare no competing financial interests.

## Funding

MP was supported by Agriculture and Food Research Initiative Competitive Grant 2013-6715-20966 from the U.S. Department of Agriculture to Patrick Burns.

## References

[1] Li MO, Wan YY, Sanjabi S, Robertson A-KL, Flavell RA. TRANSFORMING GROWTH FACTOR-ß REGULATION OF IMMUNE RESPONSES. Annu Rev Immunol. 2006 Apr;24(1):99–146. (http://dx.doi.org/10.1146/annurev.immunol.24.021605.090737)

[2] Theoharides TC, Alysandratos K-D, Angelidou A, Delivanis D-A, Sismanopoulos N, Zhang B, et al. Mast cells and inflammation. BiochimBiophys Acta. 2012 Jan;1822(1):21–33. (http://dx.doi.org/10.1016/j.bbadis.2010.12.014)

[3] Galli SJ, Tsai M. IgE and mast cells in allergic disease. Nature Medicine. 2012 May 4;18(5):693–704. (http://dx.doi.org/10.1038/nm.2755)

[4] Saluja R, Ketelaar ME, Hawro T, Church MK, Maurer M, Nawijn MC. The role of the IL-33/IL-1RL1 axis in mast cell and basophil activation in allergic disorders. Molecular Immunology. 2015 Jan;63(1):80–5. (http://dx.doi.org/10.1016/j.molimm.2014.06.018)

[5] Polukort SH, Rovatti J, Carlson L, Thompson C, Ser-Dolansky J, Kinney SRM, et al. IL-10 Enhances IgE-Mediated Mast Cell Responses and Is Essential for the Development of Experimental Food Allergy in IL-10–Deficient Mice. J Immunol. 2016 May 6;196(12):4865–76. (http://dx.doi.org/10.4049/jimmunol.1600066)

[6] Qayum AA, Paranjape A, Abebayehu D, Kolawole EM, Haque TT, McLeod JJA, et al. IL-10–Induced miR-155 Targets SOCS1 To Enhance IgE-Mediated Mast Cell Function. J Immunol. 2016 Apr 29;196(11):67–67. (http://dx.doi.org/10.4049/jimmunol.1502240)

[7] Ganeshan K, Johnston LK, Bryce PJ. TGF‐ 1 Limits the Onset of Innate Lung Inflammation by Promoting Mast Cell-Derived IL-6. J Immunol. 2013 Apr 29;190(11):8–8. (http://dx.doi.org/10.4049/jimmunol.1203362)

[8] Ganeshan K, Bryce PJ. Regulatory T Cells Enhance Mast Cell Production of IL-6 via Surface-Bound TGF‐. J Immunol. 2011 Dec 9;188(2):603–603. (http://dx.doi.org/10.4049/jimmunol.1102389)

[9] Tsukamoto H, Nishikata R, Senju S, Nishimura Y. Myeloid-Derived Suppressor Cells Attenuate TH1 Development through IL-6 Production to Promote Tumor Progression. Cancer Immunology Research. 2013 Apr 29;1(1):76–76. (http://dx.doi.org/10.1158/2326-6066.CIR-13-0030)

[10] Morales JK, Saleem SJ, Martin RK, Saunders BL, Barnstein BO, Faber TW, et al. Myeloid-derived suppressor cells enhance IgE-mediated mast cell responses. J Leukoc Biol. 2013 Dec 12;95(4):50–50. (http://dx.doi.org/10.1189/jlb.0913510)

[11] Zhu J. T helper 2 (Th2) cell differentiation, type 2 innate lymphoid cell (ILC2) development and regulation of interleukin-4 (IL-4) and IL-13 production. Cytokine. 2015 Sep;75(1):24–24. (http://dx.doi.org/10.1016/j.cyto.2015.05.010)

[12] Gould HJ, Sutton BJ. IgE in allergy and asthma today. Nature Reviews Immunology. 2008 Mar;8(3):17–17. (http://dx.doi.org/10.1038/nri2273)

[13] Greer AM, Wu N, Putnam AL, Woodruff PG, Wolters P, Kinet J-P, et al. Serum IgE clearance is facilitated by human FcϵRI internalization. J Clin Invest. 2014 Feb 24;124(3):98–98. (http://dx.doi.org/10.1172/JCI68964)

[14] Yamazaki S, Nakano N, Honjo A, Hara M, Maeda K, Nishiyama C, et al. The Transcription Factor Ehf Is Involved in TGF-β–Induced Suppression of FcϵRI and c-Kit Expression and FcϵRI-Mediated Activation in Mast Cells. J Immunol. 2015 Aug 21;195(7):35–35. (http://dx.doi.org/10.4049/jimmunol.1402856)

[15] Kubiczkova L, Sedlarikova L, Hajek R, Sevcikova S. TGF-β – an excellent servant but a bad master. J Transl Med. 2012;10(1):183. (http://dx.doi.org/10.1186/1479-5876-10-183)

